# An ultrasensitive fiveplex activity assay for cellular kinases

**DOI:** 10.1101/687715

**Authors:** Christian M. Smolko, Kevin A. Janes

**Affiliations:** Department of Biomedical Engineering, University of Virginia, Charlottesville, VA 22908; Department of Biochemistry & Molecular Genetics, University of Virginia, Charlottesville, VA 22908

## Abstract

Protein kinases are enzymes whose abundance, protein-protein interactions, and posttranslational modifications together determine net signaling activity in cells. Large-scale data on cellular kinase activity are limited, because existing assays are cumbersome, poorly sensitive, low throughput, and restricted to measuring one kinase at a time. Here, we surmount the conventional hurdles of activity measurement with a multiplexing approach that leverages the selectivity of individual kinase-substrate pairs. We demonstrate proof of concept by designing an assay that jointly measures activity of five pleiotropic signaling kinases: Akt, IκB kinase (IKK), c-jun N-terminal kinase (JNK), mitogen-activated protein kinase (MAPK)-extracellular regulated kinase kinase (MEK), and MAPK-activated protein kinase-2 (MK2). The assay operates in a 96-well format and specifically measures endogenous kinase activation with coefficients of variation less than 20%. Multiplex tracking of kinase-substrate pairs reduces input requirements by 25-fold, with ~75 μg of cellular extract sufficient for fiveplex activity profiling. We applied the assay to monitor kinase signaling during coxsackievirus B3 infection of two different host-cell types and identified multiple differences in pathway dynamics and coordination that warrant future study. Because the Akt–IKK–JNK–MEK–MK2 pathways regulate many important cellular functions, the fiveplex assay should find applications in inflammation, environmental-stress, and cancer research.

## INTRODUCTION

Phosphorylation is a major posttranslational modification of proteins in cells^1^. Regulated protein phosphorylation is important for signal transduction and requires the catalytic activity of protein kinases^2^. Overall enzymatic activity is controlled at multiple levels and time scales depending on the cellular kinase^3^. Activation or inhibition can be achieved by kinase phosphorylation [e.g., mitogen-activated protein kinases (MAPKs), glycogen synthase kinase-3 (GSK3)] or allosteric protein-protein interactions [cyclin-dependent kinases (CDKs), protein kinase A (PKA)]. For some kinases, net cellular activity is altered by protein stabilization [nuclear factor-κB-inducing kinase (NIK)] or transcriptional upregulation [proviral integrations of moloney (PIM) kinases]. Mechanisms of kinase regulation are not mutually exclusive; for example, IκB kinase (IKK) is a holoenzyme complex controlled bidirectionally through phosphorylation and protein-protein interactions^4^. Reliably tracking cellular kinase signaling at scale remains highly pertinent to systems-level investigations of the biological response to complex perturbations^5–8^.

Facets of kinase regulation can be captured by modern -omics approaches^9^, but compared to direct measures of kinase catalytic activity, the information is incomplete even for well-studied pathways. A kinase-substrate pair recognized for its specificity is the Thr-Tyr bisphosphorylation of extracellular-regulated kinase (ERK) by MAPK-ERK kinase (MEK)^10^. High-throughput monitoring of ERK phosphorylation has been standardized with reverse-phase protein arrays using validated antibodies^11^. However, ERK can be dephosphorylated at Thr or Tyr by cellular phosphatases^12,13^, yielding mono-phosphorylated species with measurable activity^14^ but altered antibody epitopes^15^. Phosphorylation-activity discrepancies may be considerable given reports that ERK is mostly mono-phosphorylated as soon as ~30 min after growth-factor or cytokine stimulation^16,17^. Full activation of some kinases, such as Akt and c-jun N-terminal kinase (JNK), requires separate phosphorylation events that are catalyzed specifically^18,19^ or preferentially^20^ by different upstream kinases. Decoding the effect of such combinatorial modifications is a long-standing challenge^21,22^. Other kinase domains coexist as separate transcript variants with different regulatory and effector functions. For example, MAPK-activated protein kinase-2 (MK2) is normally expressed as two isoforms in roughly equal abundance, but only the long isoform is bound to its upstream activating kinase, p38 (Ref. ^23,24^). MK2–p38 complexes show enhanced activity toward specific protein substrates^25^. Certain phosphorylation sites may sometimes serve as acceptable substitutes for cellular kinase activity, yet there remains a need for modern platforms that quantify panels of activity directly and specifically.

Mass spectrometry has quantified *in vitro* phosphorylation of peptide substrates in whole cell extracts^26^, but unfractionated assays cannot attribute activity to individual kinases^27^. Specificity is provided by immune complex kinase assays, which remain little changed since they were introduced four decades ago^28^. In the standard assay, endogenous kinase immunoprecipitates are incubated with recombinant substrate and [γ-^32^P]ATP. The mixture is separated by polyacrylamide gel electrophoresis, and substrate phosphorylation is detected by autoradiography. We and others previously improved assay throughput by microplate immunoprecipitation and end-product isolation on 96-well phosphocellulose filters^29,30^. Sensitivity remained poor for most kinases, however, requiring hundreds of micrograms of cell extract per kinase reaction due to the radiotracer requiring operating conditions at or below the K_m,ATP_ for each enzyme^31^. More-fundamental innovations in target capture, enzymology, and detection are required for high-throughput kinase activity assays to compare favorably with competing alternatives.

One untapped opportunity lies in the phylogeny of protein kinases with respect to their substrate requirements^32^. The binding cleft of most kinase domains recognizes specific flanking residues at amino-acid positions surrounding the phosphoacceptor^33^. Some kinases possess additional docking interactions that enhance specific activity toward *bona fide* substrates^34^. Proteins lacking the proper docking sites or flanking residues for a kinase will not be phosphorylated efficiently or at all. Thus, shrewd mixtures of nonoverlapping kinase-substrate pairs could theoretically react together as a pool, provided that there were no K_m,ATP_ limitations and phosphorylated substrates could be deconvolved at the end. Analogous pooling-and-deconvolution strategies have been demonstrated with barcoded cancer cell lines^35^, but such an approach has not been considered for *in vitro* biochemistry.

In this work, we illustrate the potential for kinase-substrate multiplexing by developing a method that immunopurifies at least five endogenous kinases from different subfamilies with nonoverlapping substrate specificity. The pooled cellular immunoprecipitate catalyzes phosphorylation of five cognate substrates engineered with distinct epitope tags to enable deconvolution and quantitation on barcoded anti-tag microspheres. We actualize the concept of multiplex activity profiling for five phylogenetically diverse protein kinases—Akt, IKK, JNK, MEK, and MK2—that are broadly implicated in cellular regulation. We confirm kinase-substrate specificity both pharmacologically and by omitting individual kinases from the immunoprecipitation. Assay sensitivity approaches that of immunoblots^36^, which are widely used but can only quantify the abundance of individual phosphoproteins or kinase posttranslational modifications. Using the assay, we tracked pathogen-induced signaling dynamics in two different host cells for the positive-strand RNA virus, coxsackievirus B3 (CVB3), measuring 320 kinase activities in one day. This fiveplex panel should be similarly useful for deciphering kinase activation in response to growth factors, cytokines, and environmental stresses relevant to inflammation and cancer.

## RESULTS

### Assay design for multiplex kinase activity profiling in a 96-well format

To perform high-throughput immunoprecipitations, we begin with 96-well microplates conjugated to a Protein A/G chimera that binds the Fc domain of most IgG species. Antibody binding capacity of each microwell is ~0.5 μg, which is split empirically among the anti-kinase antibodies to be used in the multiplex immunoprecipitation (considerations for the antibody mixture are described below). Nondenatured cell lysates are incubated with antibody-bound plates and washed at low stringency to preserve kinase-associated regulators as a co-immunoprecipitate (Supplemental Fig. S1). Each immunopurified kinase requires an inert protein substrate that is phosphorylated specifically and efficiently by the kinase at a known residue. The substrate is cloned and purified as a recombinant protein harboring a 3x epitope tag unique to that substrate (Fig. 1A and Methods). Mixtures of kinase-specific substrates are added to the plate-bound immunoprecipitate along with physiological concentrations of ATP (1 mM) to maximize catalysis. Substrate concentrations are kept low (< 20 pM) for high-sensitivity detection (see below). Reactant depletion is avoided by starting with minimal input lysate (≤100 μg) and by down-titrating antibody amounts for kinases that are very abundant, catalytically active, or efficiently immunoprecipitated. The dynamic range of the *in vitro* reaction is large enough that reoptimization for different biological settings is minimal or unnecessary (see below).

**Figure 1.**
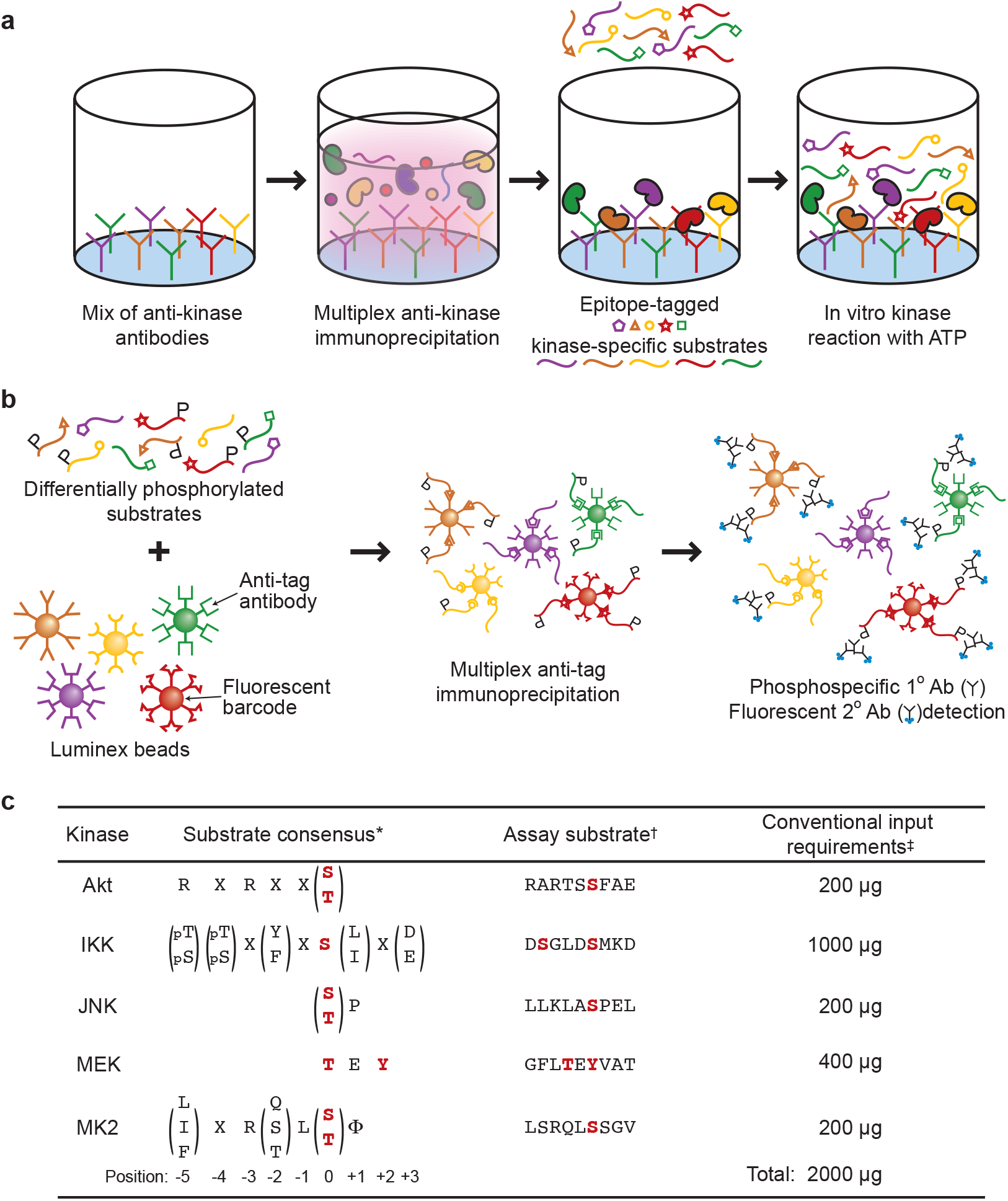
A two-step multikinase activity assay combining pooled immunoaffinity purification and phosphosubstrate deconvolution. **(a)** Titrated mixtures of anti-kinase antibodies are Fc-immobilized on Protein A/G-functionalized microplates to immunoprecipitate a target pool of endogenous kinases. Catalytic activity of the immunoprecipitated kinases is measured by incubating with ATP and kinase-specific recombinant substrates harboring separate epitope tags. After the *in vitro* kinase reaction, substrates are separated on magnetic beads that are surface conjugated with an anti-tag antibody and dyed with a specific fluorescent barcode. The extent of substrate phosphorylation is quantified by indirect immunofluorescence using antibodies specific to the kinase-catalyzed phosphoepitope. **(c)** Protein kinases selected in the original fiveplex design are shown alongside their published substrate consensus sequences, assay substrates, and lysate input requirements when measured by conventional immune complex kinase assays. *Consensus sequences for kinase substrates as defined by Obata et al.^97^, Hutti et al.^98^, Dérijard et al.^99^, Seger et al.^10^, and Manke et al.^100^. ^†^Further substrate details are available in Table 1. ‡Representative input requirements from Hill and Hemmings^101^, Mihalas and Meffert ^102^, Whitmarsh and Davis^103^, Okano and Rustgi^104^, and Janes et al.^30^.

Terminated kinase reactions are deconvolved and quantified by bead-based sandwich immunoassay using a Luminex instrument (Fig. 1B). For substrate capture, mouse anti-epitope tag antibodies are separately conjugated to the surface of microspheres fluorescently barcoded to indicate epitope-tag identity. The separate epitope-capture beads are pooled and incubated with the terminated kinase reaction in 96-well plates to demultiplex the solution. The extent of phosphorylation on each bead is then detected with phosphospecific rabbit antibodies recognizing the residue on the substrate modified by the cognate kinase. Phosphospecific immunoreactivity is quantified with anti-rabbit secondary labeled with phycoerythrin, which is scanned along with the fluorescent barcode of each bead in the mixture. Overall, the two-step procedure creates a high-throughput, nonradioactive assay for specifically profiling the activity of multiple kinases together.

**Table 1.** Antibody (Ab), substrate, and detection conditions for the fiveplex kinase activity assay. CST, Cell Signaling Technology; SCBT, Santa Cruz Biotechnology; R&D, R&D Systems; MA015, ADI-KAP-MA015.

| Kinase | Immunoprecipitating Ab |  | Substrate |  |  | Detection Ab |  |
| --- | --- | --- | --- | --- | --- | --- | --- |
|  | Source | Amount | 3x Tag | Protein | Amount | Source | Dilution |
| Akt | Millipore #05-591 | 20 ng | VSVG | GSK3α(1-97) | 0.1 μg | CST #9316 | 1:5000 |
| IKK | CST #2682 | 220 ng | HA | IκBα(1-62) | 0.1 μg | R&D #AF4809 | 1:5000 |
| JNK | SCBT #sc-474 | 80 ng | GluGlu | c-jun | 1 μg | CST #9164 | 1:2500 |
| MEK | Abcam #ab178876 | 80 ng | Myc | ERK2(K52R) | 0.1 μg | CST #9101 | 1:2500 |
| MK2 | Enzo #MA015 | 40 ng | FLAG | HSP27 | 1 μg | CST #2401 | 1:200 |

### Optimization of a fiveplex Akt–IKK–JNK–MEK–MK2 activity assay

To realize the design concept, we selected five pleiotropic protein kinases with complex modes of regulation (see Introduction) and distinct substrate preferences (Fig. 1C). Akt is a basophilic kinase involved in cell survival and metabolism^37^. IKK is a multikinase signalsome that acts as a central activator of inflammation through the nuclear factor-κB transcription factor^38^. JNK is a stress-activated MAPK that broadly regulates gene expression and various cytoplasmic processes^39^. MEK is a proliferative MAPK kinase activated by growth factor receptors as well as many oncogenes^40^. MK2 is a downstream effector of p38 MAPK signaling and an important posttranscriptional regulator of mRNA stability^41^. These five kinases are convergence points for many upstream activators and downstream cellular effector pathways^42^, providing strong justification for measuring them as a panel.

Akt–IKK–JNK–MEK–MK2 are potently activated together by the combined stimulation of cells with tumor necrosis factor (TNF), epidermal growth factor (EGF), and high concentrations of insulin^43–45^. This T+E+I condition was used as a positive control when optimizing the individual anti-kinase titrations and input lysate for the multiplex immunoprecipitation (Fig. 1A). Detecting specific phosphoepitopes (Fig. 1B) enabled use of full-length substrates; those with enzymatic activity were rendered inert by truncation [GSK3α(1-97)] or mutation [ERK2(K52R)]. For substrates purified as glutathione S-transferase (GST) fusion proteins (see Methods), the GST domain was cleaved off to avoid inadvertent heterodimer formation between substrates^46^. Considering the number of epitope-capture beads measured by the Luminex instrument, the maximum theoretical binding capacity per reaction was ~0.5 fmol substrate (Supplementary Note). Using 0.1–1 μg of each substrate per reaction ensured saturated binding of the bead surface while remaining >1000-fold below the K_M_ of the reaction to avoid diluting out the stoichiometry of phosphorylation. We optimized an effective V_max_ for each reaction by titrating the amount of immunoprecipitating antibody and input lysate until substrate depletion was negligible for all kinases in the positive-control lysate. Optimal conditions were achieved with 75 μg cell extract, or ~375,000 cells^47^. The cellular input is 25-fold less than would be required to measure Akt–IKK–JNK–MEK–MK2 activity by conventional assays (Fig. 1C).

### Linear dynamic range of the fiveplex kinase activity assay is sufficient for high-throughput profiling

We determined signal to noise of the assay by measuring 15 technical replicates of extracts prepared from HT29 colon adenocarcinoma cells with or without T+E+I stimulation. The Luminex xMAP instruments report median phycoerythrin intensity of 200–400 beads from each barcode, which served as the relative fluorescence unit (RFU) for kinase activity. Background staining was assessed from *in vitro* reactions with blank immunoprecipitations containing lysis buffer alone. For all kinases except for MEK, the RFU for blank samples was below the 95% confidence interval of unstimulated extracts, suggesting measurable baseline activity (Fig. 2A–E). Negligible MEK catalysis in quiescent cells was likely the result of cytoplasmic phosphatases, which deactivate MEK under resting conditions^48^. Stimulation with T+E+I for 15 min increased the assay RFUs by ~8–30-fold, and the background-corrected coefficient of variation for all measured kinases was less than 20% (Fig. 2F). Using the unstimulated and T+E+I-stimulated samples as negative and positive controls, we calculated separate Z’ factors for each kinase assay in the multiplex pool^49^. Z’ factors were all greater than 0.5 and most were greater than 0.7, indicating a separation band between controls that is excellent for high-throughput profiling^50^.

**Figure 2.**
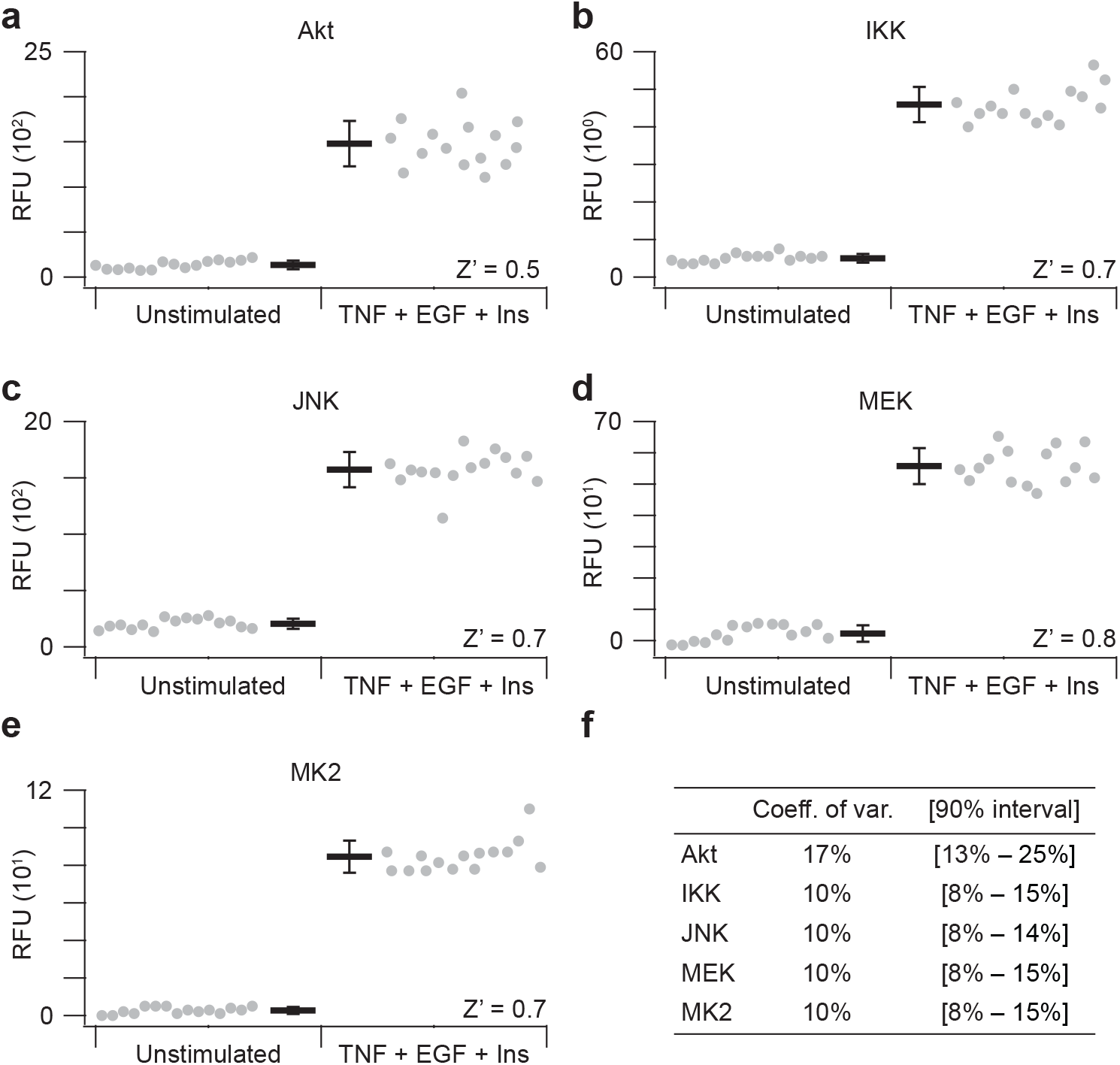
Positive- and negative-control extracts are reproducibly separated in the fiveplex kinase activity assay. Substrate phosphorylation was measured in parallel for **(a)** Akt activity toward 3xVSVG-GSK3α(1-97) on Ser21, **(b)** IKK activity toward 3xHA-IκBα(1-62) on Ser32+Ser36, **(c)** JNK activity toward 3xGluGlu-c-jun on Ser73, **(d)** MEK activity toward 3xMyc-ERK2(K52R) on Thr202+Tyr204, and **(e)** MK2 activity toward 3xFLAG-HSP27 on Ser82. Data are shown as the mean relative fluorescence unit (RFU) ± s.d. of *n* = 15 assay replicates of 75 μg extract from unstimulated HT29 cells (left) or after treatment with 20 ng/ml TNF, 100 ng/ml EGF, and 500 ng/ml insulin for 15 min (TNF + EGF + Ins; right). Z’ factors for the band separation of each kinase assay are shown according to Zhang et al.^49^. **(f)** Assay coefficients of variation for the TNF + EGF + Ins positive control extracts. Confidence intervals are estimated by McKay’s transformation^105^.

We next examined the linear dynamic range of the assay by varying the proportion of T+E+I-stimulated and unstimulated extract in 75 μg total input lysate. Keeping the amount of cellular extract constant avoids concentration differences that might confound the immunoprecipitation. From 0% to 100% stimulated extract, all five kinase activities were linear with R^2^ values all greater than 0.96 for HT29 cell extracts (Fig. 3A–E), indicating lack of substrate depletion under the assay conditions. Linear response characteristics obviate the need for logistic curve fitting of standards to make quantitative measurements^51^. We repeated the titration with extracts from AC16-CAR cells, a human ventricular cardiomyocyte cell line^52^ engineered to express the coxsackievirus and adenovirus receptor^13^. When using AC16-CAR extracts, we again observed linear behavior for all five kinases in with R^2^ values of 0.94 or better (Supplemental Fig. S2). For JNK, a hyperbolic response curve was slightly favored over a linear model (*p* ~ 0.025 by χ^2^ goodness-of-fit test), suggesting substrate depletion is mild for highly kinase-active cells, such as AC16-CAR (see below). Considering the differences in lineage [mesodermal (AC16-CAR) vs. endodermal (HT29)] and basal signaling (see below), the titration results suggest that the assay will accurately quantify kinase activities in diverse cell types. If substrate depletion is extensive, revised assay conditions should increase the amount of substrate or decrease the amount of immunoprecipitating antibody (see Methods).

**Figure 3.**
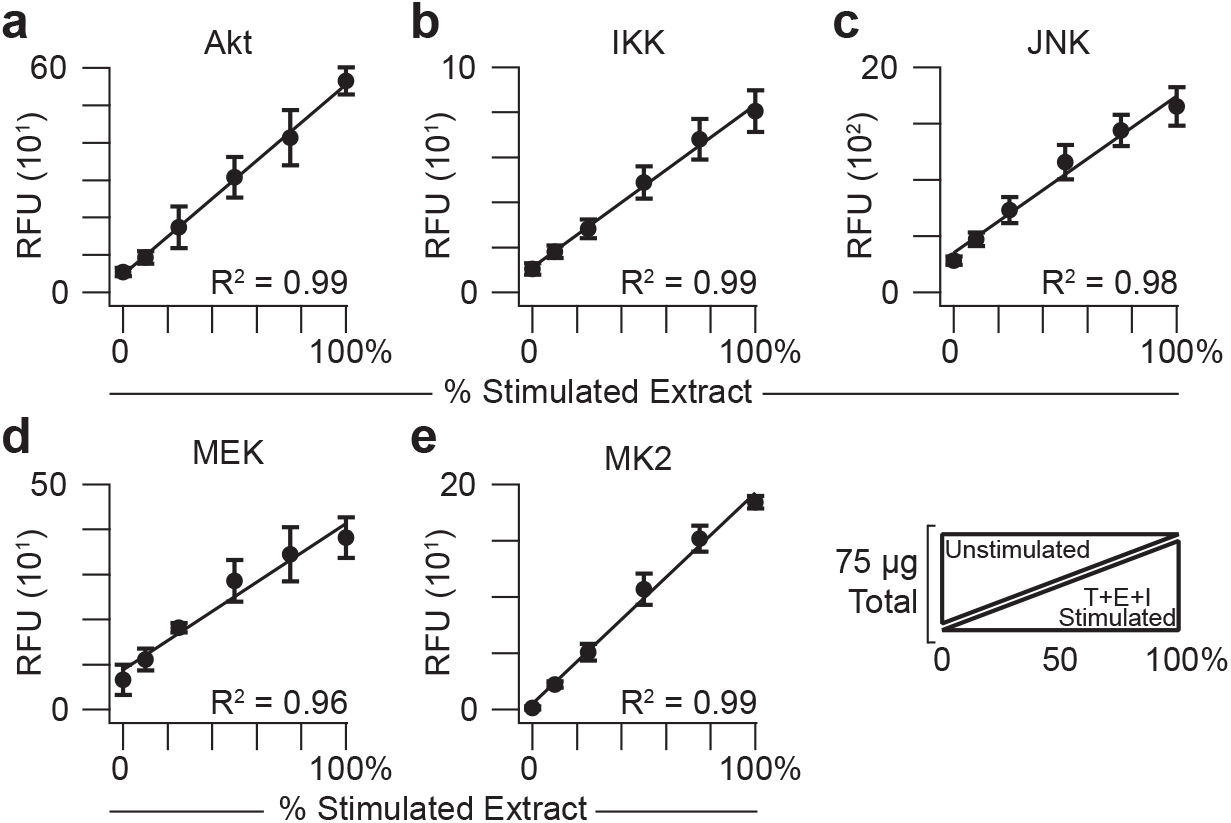
The fiveplex kinase assay quantifies endogenous catalytic activity linearly. Relative *in vitro* phosphorylation of substrates for **(a)** Akt, **(b)** IKK, **(c)** JNK, **(d)** MEK, and **(e)** MK2 was measured together as a function of proportionately stimulated cell extract added. Data are shown as the mean relative fluorescence unit (RFU) ± s.d. of *n* = 4 assay replicates of 75 μg extract mixed in the indicated proportions of unstimulated HT29 cells and cells treated with 20 ng/ml TNF, 100 ng/ml EGF, and 500 ng/ml insulin for 15 min (T+E+I stimulated).

### Kinase-substrate specificity is retained during multiplexing

Kinase-substrate pairs were selected to have zero overlap (Fig. 1C), but multiplexing could give rise to new confounding variables that prevent the *in vitro* reactions from proceeding independently. We tested the specificity of the multiplex assay by systematically substituting naïve IgG for one immunoprecipitation antibody at the start of the procedure. With each kinase, the removal of its immunoprecipitation antibody reduced the measured activity to background levels (Fig. 4A–E, arrows). Moreover, removal of one immunoprecipitation antibody had minimal impact on the remaining four kinase activities in the assay. The data suggest that proteins co-immunoprecipitating with a kinase (e.g., Supplementary Fig. S1) do not have a discernible influence on the activity of other kinases in the fiveplex panel.

**Figure 4.**
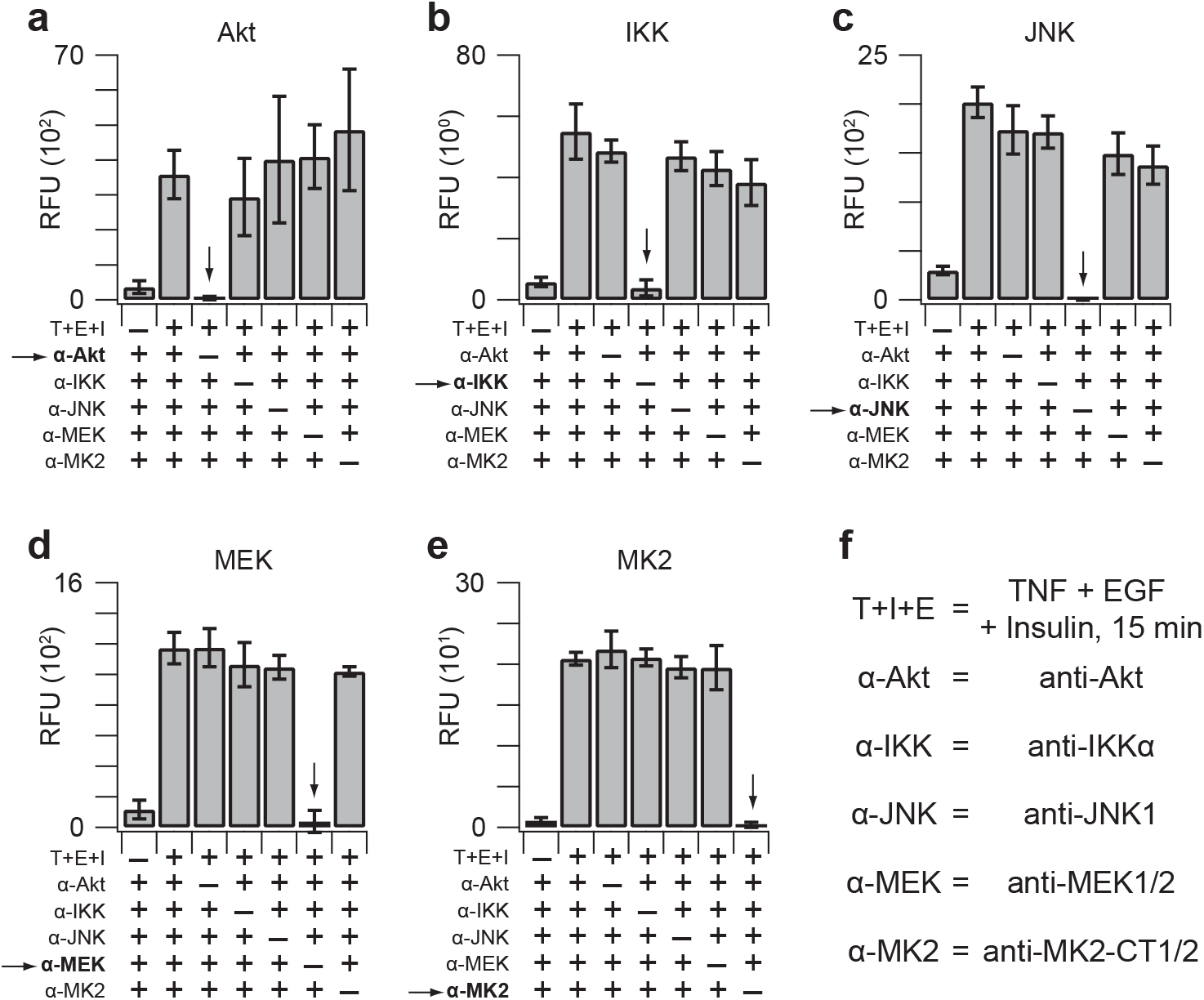
Kinase-specific substrate phosphorylation requires kinase-specific immunoprecipitations. Leave-one-out immunoprecipitations were measured using 75 μg extract from HT29 cells stimulated with 20 ng/ml TNF, 100 ng/ml EGF, and 500 ng/ml insulin for 15 min (T+E+I). Relative *in vitro* phosphorylation of substrates for **(a)** Akt, **(b)** IKK, **(c)** JNK, **(d)** MEK, and **(e)** MK2 was measured with the fiveplex kinase assay. Data are shown as the mean relative fluorescence unit (RFU) ± s.d. of *n* = 4 assay replicates. **(f)** Isoform-specific information about the immunoprecipitating antibodies. The MK2 antibody recognizes both isoforms with different C termini (CT).

To perturb catalytic activity specifically, we pharmacologically inhibited each kinase separately and measured the assay response (Fig. 5A–E). Small-molecule inhibitors were used differently in the assay depending on their mechanism of action (Fig. 5F). Reversible inhibitors of upstream enzymes (PI3Ki, p38α/βi), allosteric inhibitors of activation (MEKi), and covalent inhibitors (JNKi) were added to cultures before T+E+I stimulation to inhibit kinase activity *in cellulo*. A potent *in cellulo* inhibitor for IKK could not be identified; therefore, we inhibited IKK directly by using a reversible ATP-competitive small molecule of one subunit (IKKβi) together with 100-fold less ATP during the *in vitro* kinase reaction (see Methods). For Akt, JNK, MEK, and MK2, measured kinase activity was abolished upon pathway inhibition in cells (Fig. 5A,C–E). *In vitro* IKK inhibition reduced its measured activity by ~50% (Fig. 5B), consistent with this small molecule inhibiting one of the two catalytic subunits of the IKK signalsome, both of which phosphorylate the substrate^53–55^. IKKβi had little-to-no effect on the other kinases *in vitro*. However, the accompanying reduction in ATP lowered the overall extent of substrate phosphorylation by 4–30-fold and rendered MEK so enzymatically deficient that it could not be detectably activated (Fig. 5A,C–E; right). Such working ATP concentrations are typical in conventional radioassays, emphasizing the importance of using physiological ATP concentrations to measure kinase activity with maximum sensitivity.

**Figure 5.**
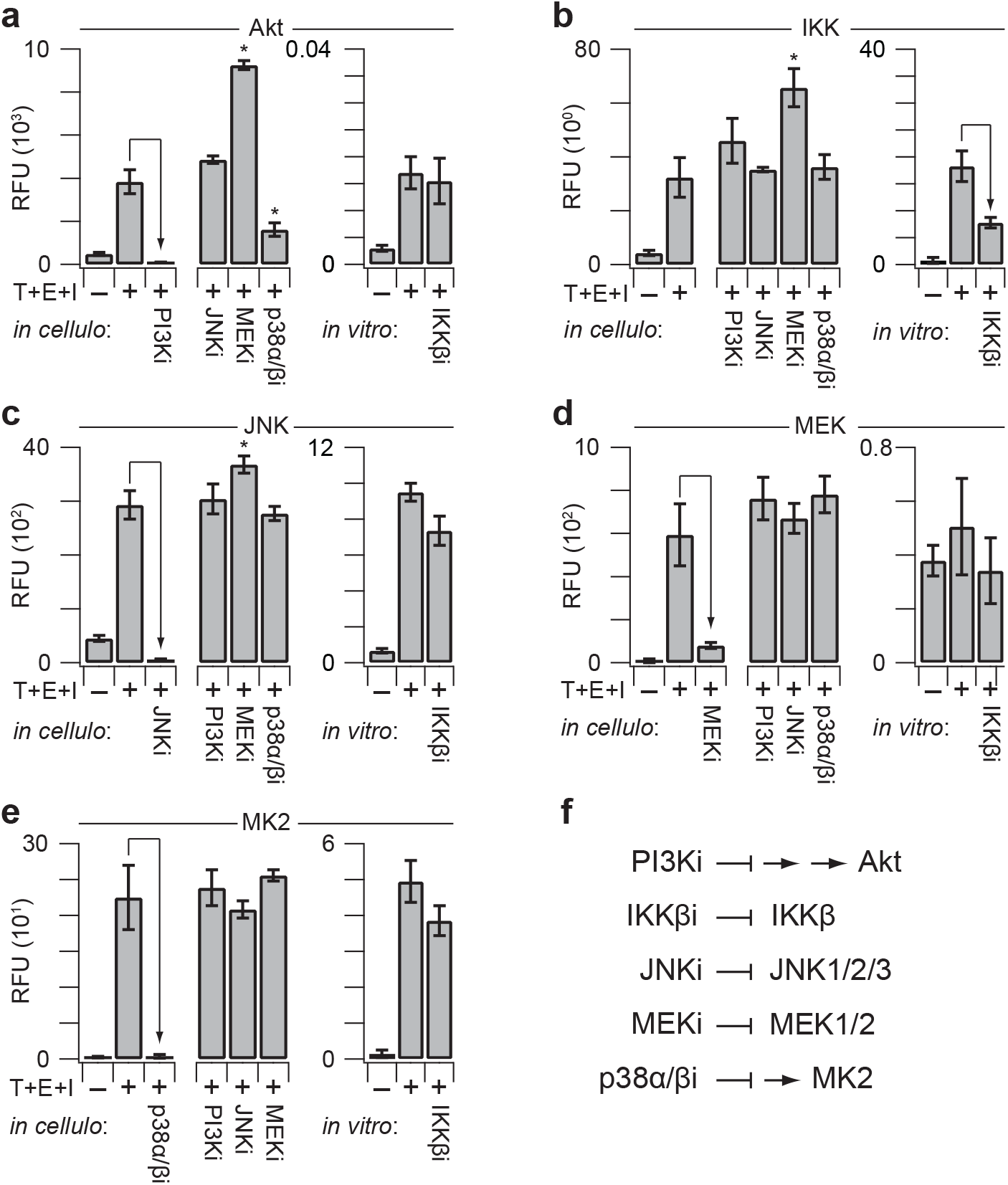
The fiveplex kinase assay detects loss of activity along with secondary cellular adaptations to small-molecule inhibitors. HT29 cells were pretreated with 50 μM LY294002 for 1 hr (PI3Ki), 1 μM JNK-IN-8 for 3 hr (JNKi), 100 nM trametinib for 2 hr (MEKi), or 100 μM SB202190 for 1 hr (p38α/βi) and then stimulated with 20 ng/ml TNF, 100 ng/ml EGF, and 500 ng/ml insulin for 15 min (T+E+I). For *in vitro* inhibition, immunoprecipitates of T+E+I-stimulated extracts were incubated with 10 μM SC-514 (IKKβi) during the kinase reaction, which contained reduced ATP (see Methods). Relative *in vitro* phosphorylation of substrates for **(a)** Akt, **(b)** IKK, **(c)** JNK, **(d)** MEK, and **(e)** MK2 was measured with the fiveplex kinase assay. Data are shown as the mean relative fluorescence unit (RFU) ± s.d. of *n* = 4 assay replicates. \**p* < 0.01 by ANOVA and Tukey’s posthoc test for significant secondary *in cellulo* adaptations compared to the corresponding T+E+I positive control. **(f)** Abbreviated mechanisms of action for the small-molecule inhibitors used *in cellulo* and *in vitro*. Upstream kinases are abstracted for clarity.

Inhibiting kinases *in cellulo* will lead to secondary changes downstream of the primary target, and we observed several alterations consistent with the literature. MEKi caused an increase in T+E+I-stimulated Akt activity (Fig. 5A) as well as JNK activity (Fig. 5C), likely due to loss of feedback inhibition through receptor tyrosine kinases^56^ and dual-specificity phosphatases^57^ respectively. T+E+I-stimulated activation of IKK was also enhanced after MEKi pretreatment (Fig. 5B), an effect that is probably secondary to the enhanced activation of Akt, which can augment IKK signaling in some settings^58^. Conversely, Akt activity was inhibited when the MK2 pathway was targeted *in cellulo*, possibly reflecting alterations in phospholipid levels or off-target effects of p38α/βi on Akt directly^59,60^. Overall, secondary and off-target effects were infrequent and uncoupled from the results of the leave-one-out immunoprecipitations (Fig. 4), supporting the specificity of the assay to deconvolve individual kinase activities.

### Akt–IKK–JNK–MEK–MK2 activity profiling of host-cell signaling dynamics during CVB3 infection

Akt–IKK–JNK–MEK–MK2 transduce signaling from many environmental stimuli, including pathogens^61,62^. As a representative cellular perturbation to examine with the fiveplex assay, we selected acute infection by the picornavirus CVB3. During transmission and pathogenesis, CVB3 infects multiple cell types^63^, including the gut epithelium (e.g., HT29 cells^64^) and cardiomyocytes (e.g., AC16-CAR cells^13^). In other model cell systems, CVB3 impacts all five kinases in the assay^65–68^. For one cardiomyocyte cell line (HL-1), time-dependent CVB3-induced changes in 8–9 host-cell phosphoproteins were monitored longitudinally by commercial phospho-ELISA measurements of cell extracts^69,70^. We recognized the opportunity to appraise similarities and differences between CVB3-infected cell types, as well as between kinase catalytic activity and phosphoprotein surrogates.

HL-1 cardiomyocytes require a proprietary culture medium with many supplements^71^ that we feared would corrupt meaningful comparisons in baseline activity with HT29 cells. Therefore, we substituted AC16-CAR cardiomyocytes, infecting them and HT29 cells at the same plating density with a high CVB3 multiplicity of infection (MOI = 10). Cells were lysed at multiple time points from 0–8 hrs to capture the signaling dynamics from CVB3 internalization until cytopathic effect and cell death. Extracts were prepared in biological quadruplicate and measured all together to remove any possibility of batch effects. The study required 320 kinase activity measurements (2 cell lines × 8 time points × 4 biological replicates × 5 kinases), which were completed in one day with the multiplex assay (Fig. 6A–E).

**Figure 6.**
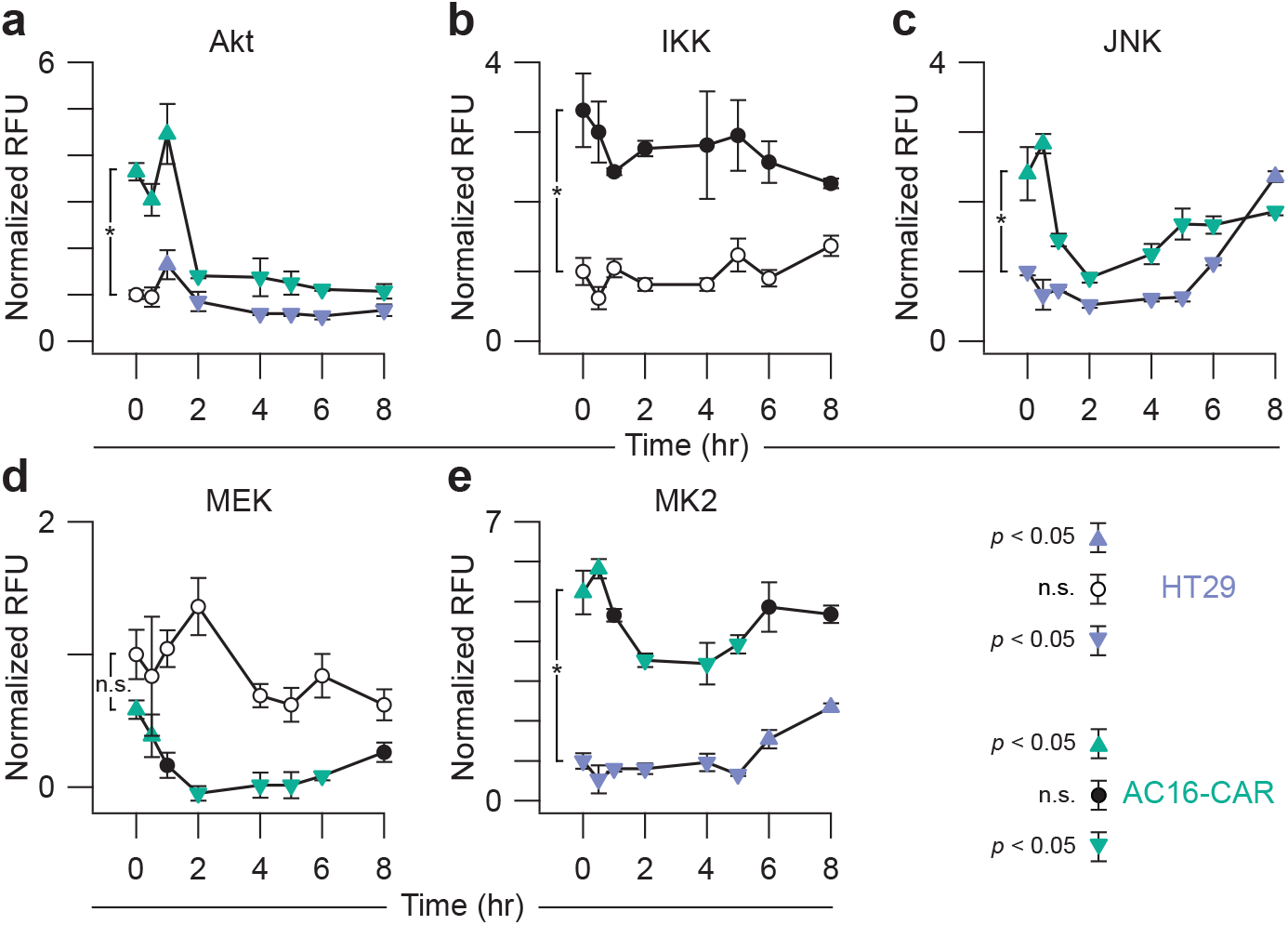
Multiplex kinase-activity profiling of time-dependent and cell-specific responses to coxsackievirus B3 (CVB3) infection. HT29 and AC16-CAR cells were infected with CVB3 at a multiplicity of infection of 10 at time = 0 hr. From extracts collected at each time point, relative *in vitro* phosphorylation of substrates for **(a)** Akt, **(b)** IKK, **(c)** JNK, **(d)** MEK, and **(e)** MK2 was measured with the fiveplex kinase assay. Data are shown as the mean relative fluorescence unit (RFU) ± s.d. of *n* = 4 biological replicates. \**p* < 0.05 by Student’s unpaired *t* test for differences between the HT29 and AC16-CAR 0-hr time points. Time points significantly greater than or less than other time points (*p* < 0.05 by ANOVA and Tukey’s post hoc test) are shown as upward and downward arrowheads respectively.

Before CVB3 infection, we noted several differences in baseline kinase activity per μg extract between AC16-CAR cells and HT29 cells. Despite harboring an oncogenic *PIK3CA*^*P449T*^ mutation^72^, basal Akt activity in HT29 cells was substantially lower than in AC16-CAR cells (Fig. 6A). The AC16 line is immortalized with SV40 tumor antigen, the small splice variant of which inhibits PP2A phosphatase^73^. PP2A removes activation loop phosphorylation of Akt on Thr308^74^, deactivates IKK^75^, and negatively regulates the JNK and MK2 pathways^76,77^, explaining their similarly elevated basal activities in AC16-CAR cells (Fig. 6B,C,E). PP2A inhibition with SV40 small tumor antigen also activates MEK^78^, but baseline MEK activity was comparably high in non-quiescent HT29 cells (Fig. 6D) likely because of a hyperactive *BRAF^V600E^* mutation^72^. The two lines provided considerably different initial signaling contexts to evaluate the host-cell response to CVB3 infection.

In both AC16-CAR and HT29 cells, CVB3 infection gave rise to late-phase activation of JNK and MK2 activity at 4–8 hr after infection (Fig. 6C,E). The dynamics agreed with the phosphorylation of JNK and HSP27 (an MK2 substrate) in HL-1 cardiomyocytes infected with CVB3^69,70^. MEK activity also began to rise at late times in AC16-CAR cells (Fig. 6D). Late-phase MEK activation is consistent with increased phosphorylation of ERK (a MEK substrate) in HL-1 cells at 8 hr after CVB3 infection^69,70^ and attributable to CVB3-mediated cleavage of RasGAP^68^. A similar activation was not observed or expected in HT29 cells, because MEK activity is chronically high downstream of mutant BRAF irrespective of RasGAP cleavage.

We noted the greatest deviations in activity and phosphorylation at early-to-intermediate times 0.5–4 hr after CVB3 infection. For example, loss of Akt phosphorylation on Ser473 was measured by phospho-ELISA in HL-1 cells at 16–24 hr after infection^69^, but we found that Akt activity dropped ~fourfold in AC16-CAR cardiomyocytes within 2 hr (Fig. 6A). Phosphorylation of Akt on its activation loop (Thr308) is more labile than Ser473^36^, and both phosphosites are required for full activation^79^, raising the possibility of different dephosphorylation kinetics during CVB3 infection. The phosphorylation of IκBα in HL-1 cells^69^ had little-to-no correspondence with IKK activity in AC16-CAR or HT29 cells (Fig. 6B). Phosphorylation of cellular IκBα causes its degradation^80^, and total IκBα is also cleaved independently of IKK by a CVB3-encoded protease^66^, complicating interpretation of the phospho-ELISA data^36^. In AC16-CAR cells, the multiplex kinase assay showed clear decreases in MAPK pathway signaling, with JNK, MEK, and MK2 significantly reduced within 2 hr of CVB3 infection (Fig. 6C–E). There are glimpses of pathway deactivation in JNK, ERK, and p38 phospho-ELISA data from HL-1 cells^69,70^ and immunoblot data from other CVB3 host cells^67,81^. However, the differences were small and thus ignored in prior work. By providing dynamic range to quantify both increases and decreases in cellular activity, the multiplex kinase assay is a powerful accompaniment to multi-protein data on individual phosphorylation sites.

In the CVB3-treated samples, the biological coefficient of variation in kinase signaling often exceeded technical variation (median = 26%, interquartile range: 12–38%). Replicate-to-replicate fluctuations partly reflect intrinsic noise in the timing of viral docking, internalization, endosomal escape, and logarithmic replication in the cytoplasm^82^. Accompanying host-cell adaptations should be captured in the cell extracts; if so, the multiplexed data would contain information about which kinase pathways are coupled during an adaptation. We centered the multikinase data around the mean for each cell line and time point and searched for inter-replicate fluctuations that covaried significantly (Fig. 7). Using a stringent type I error rate (α = 0.01), we found that none of the kinase-to-kinase covariations were significant in HT29 cells after multiple hypothesis-test correction, suggesting random biological noise. By contrast, there were multiple significant correlations in AC16-CAR cells. JNK and MK2 fluctuations were the strongest covariates (*R* = 0.79), further supporting that these two pathways are tightly coupled based on their activity dynamics (Fig. 6C,E). More surprising was the IKK pathway, which was highly variable in AC16-CAR cells during viral docking and the early stages of viral replication (Fig. 6B). We detected significant co-fluctuations between IKK and JNK, MEK, and MK2 (Fig. 7), raising the possibility that a common upstream activator is variably perturbed during CVB3 infection of cardiomyocytes. Although a full mechanistic elaboration is beyond the scope here, our data and analysis provide a substantive example of how multikinase activity profiling can be deployed to interrogate cellular regulation at the systems level.

**Figure 7.**
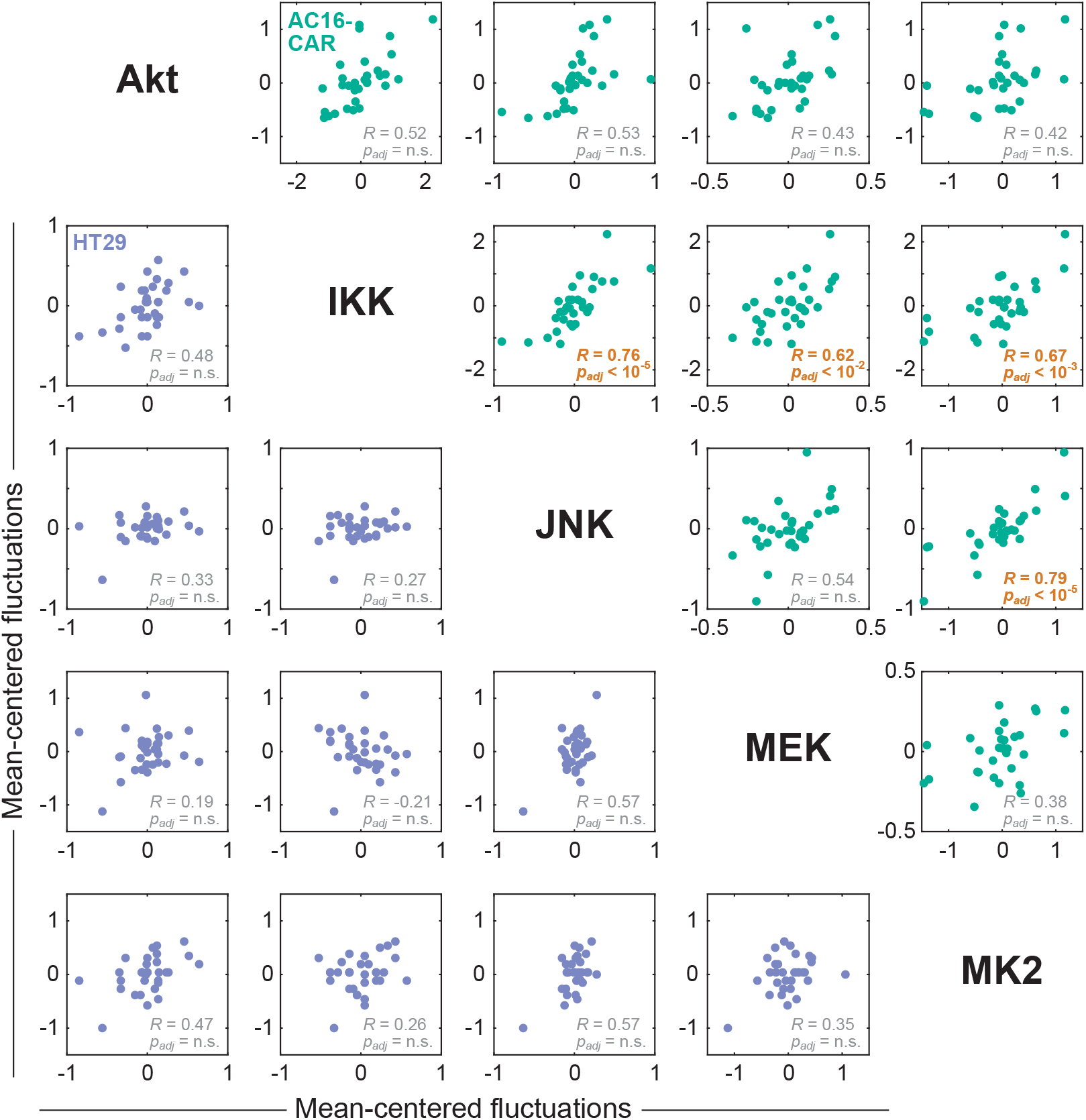
Inter-replicate fluctuation analysis of multiplex kinase activities reveals covariation between IKK and JNK–MEK–MK2 during CVB3 infection of AC16-CAR cells. Biological replicates were centered at each time point, aggregated, and assessed statistically for nonzero Pearson correlations (*R*) at 1% type I error rate, with *p* values reported after Šidák correction for multiple-hypothesis testing (*p*_*adj*_).

## DISCUSSION

The approach described here shows the substantial improvements in assay performance that are possible when multiplexing enzyme-catalyzed reactions. Tracking kinase activities in parallel reduces sample requirements and provides greater statistical power to identify coregulated signaling pathways^83^. Increases in plexity do not need to be large if throughput is not sacrificed as a tradeoff. In quantitative PCR, for example, multiplex TaqMan assays^84^ of 2– 5 targets are widely pursued as diagnostics^85^. For the activity assay (Fig. 1), the increased cost in adding a kinase is nominal once epitope-capture beads have been conjugated and recombinant substrates purified. The procedure is accessible, in that it does not require radioisotopes or specialized instrumentation beyond an xMAP bead-array reader (Luminex 100/200 analyzer, MAGPIX, or FLEXMAP 3D). Immediately, the optimized assay can be applied to studies of Akt–IKK–JNK–MEK–MK2 signaling in response to inflammation and other environmental stresses, including pathogens and activating oncogenes.

To detect multiple kinase activities, we used Luminex microspheres, recognizing their success in commercial assays that quantify panels of cellular phosphoproteins^86^. Theoretically, xMAP bead-array readers can deconvolve up to 500 analytes, raising the possibility of appending competitive or cooperative substrates to a kinase panel. For example, an assay that multiplexes ERK and GSK3 activity measurements might track specific substrates along with the phosphorylation of Myc on Thr58, a GSK3 site dependent on prior phosphorylation at Ser62 by ERK^87^. We also note that the fiveplex immunoprecipitation has not fully saturated the binding capacity of the Protein A/G microplates, with 60–100 ng of binding capacity available for a sixth nonoverlapping kinase (Table 1). The most-obvious candidate is a tyrosine kinase, which could be paired with a 3xAU1-tagged substrate. However, despite extensive attempts with c-Abl (+CrkL substrate) and JAK1/2 (+STAT1 substrate), we were unable to detect endogenous activity reliably, likely because of the autoinhibited confirmations of these kinases^88,89^. An attractive alternative are the acidophilic kinases, such as polo-like kinases, where substrates are known and regulated changes in activity have been documented^90^.

The five-kinase analysis of CVB3-infected host cells demonstrates the potential of multiplex activity profiling. Our data corroborate earlier phosphoprotein measurements of CVB3 infection in other cell contexts^69,70^ and suggest additional effects that were previously unrecognized. The AC16-CAR-specific covariation between IKK and JNK–MEK–MK2 (Fig. 7) is not apparent in the dynamic trajectories of these kinases (Fig. 6) and may reflect sample-specific adaptations to CVB3 (Ref. ^13^). One potential mechanism involves autocrine signaling circuits, which are variably time-evolving and iterative^45^. CVB3 infection of cardiomyocytes triggers autocrine TNF-family cytokines that modulate viral progeny release^70^, and several of these cytokines activate IKK–JNK–MEK–MK2 concurrently^45^. Based on RNA-seq data, AC16-CAR cells do not express *TNF* (zero transcripts per million [TPM]; M. Shah and K.A.J., unpublished observations). However, unlike HT29 cells (~1 TPM)^91^, AC16-CAR cells moderately express *TNFSF10* (~50 TPM), which declines considerably after 8 hr of CVB3 infection (~9 TPM). Furthermore, AC16-CAR cells abundantly express the cognate receptor *TNFRSF10B* (~330 TPM). TNF-related apoptosis-inducing ligand (TRAIL)-death receptor 5 (DR5) signaling through TNFSF10–TNFRSF10B will activate IKK–JNK–MEK–MK2 in many nontransformed cells^92^. The multiplex kinase-activity fluctuations are consistent with the hypothesis that autocrine TRAIL-DR5 signaling is asynchronously downregulated in cardiomyocytes during CVB3 infection.

Measurements of protein function provide information that is complementary to protein abundance or modifications on individual residues. Gaudet et al.^43^ statistically assessed the predictive information of ~10,000 cellular measurements of protein abundance, phosphorylation, cleavage, and activity. Small subsets of mixed data types were as predictive as the entire dataset, but relying on any single data type was inferior even when all of the data of that type was used^43,44^. Analogously, by analyzing tumor samples with multiple omics platforms, Hoadley et al.^93^ identified pan-cancer organizing principles that were incomplete or missing in single-platform datasets. One unfortunate consequence of a drive toward standardization is homogenization, where a handful of standardized methods are used to tackle every conceivable biological question. Cells use diverse strategies to regulate signal transduction and decision-making^94^—the approach(es) used to measure them should be just as multifaceted.

## METHODS

### Cell lysis and quantification

HT29 cells were obtained from ATCC and grown in McCoy’s 5A medium (Gibco #16600-082) plus 10% fetal bovine serum and penicillin-streptomycin. AC16 cells^52^ were purchased from Mercy Davidson and grown in DMEM/F12 medium (Gibco #11330-032) plus 12.5% fetal bovine serum and penicillin-streptomycin. Confluent plates were split 1:3 and cultured for 48 hours before use. For stimulated extracts, 20 ng/mL TNFα (PeproTech #300-01A), 100 ng/mL EGF (PeproTech #AF-100-15), and 500 ng/mL insulin (Sigma #I1882) were added to the culture media and incubated for 15 min at 37°C. For *in cellulo* inhibition, cells were pretreated with JNK-IN-8 (1 μM for 3 hr; Selleckchem #S4901), trametinib (100 nM for 2 hr; Selleckchem #S2673), SB202190 (100 μM for 1 hr; Selleckchem #S1077), or LY294002 (50 μM for 1 hr; Selleckchem #S1105). Cells were washed with ice-cold PBS and lysed in 50 mM Tris (pH 7.5) containing 50 mM β-glycerophosphate, 10 mM sodium pyrophosphate, 30 mM sodium fluoride, 1% Triton X-100, 150 mM NaCl, 1 mM benzamidine, 2 mM EGTA, 100 μM sodium orthovanadate, 1 mM dithiothreitol, 10 μg/mL aprotinin, 10 μg/mL leupeptin, 1 μg/mL pepstatin, 1 μg/mL microcystin-LR, and 1 mM phenylmethylsulfonyl fluoride (kinase lysis buffer). Clarified extracts were prepared by incubating cells with lysis buffer for 15 min on ice followed by centrifugation for 15 min at 16,162 rcf (Beckman Coulter, Microfuge® 16). Extract concentration was quantified by bicinchoninic acid assay (Pierce #23225) and frozen at –80°C until use.

### Plasmids

Bacterial plasmids expressing glutathione S-transferase (GST) fused to triple epitope tags were prepared by directionally cloning in-frame epitope tags into the BamHI and EcoRI restriction sites of pGEX-4T-1 (GE Healthcare Life Sciences #28954549). PCR amplicons were prepared from a PAGE-purified oligonucleotide template with forward primers containing a BglII restriction site and reverse primers containing HindIII–BamHI–EcoRI sites (Supplementary Table S1). GST-3xVSVG-GSK3α(1-97) was cloned into the BamHI and EcoRI sites of pGEX-4T-1 (3xVSVG) by PCR of human GSK3α(1-97) from pCMV-SPORT6 (Open Biosystems #MHS6278-202756882) with primers containing restriction sites for BamHI and EcoRI. 3xVSVG-GSK3α(1-97)-His_6_ was cloned into the NdeI and XhoI sites of pET-29b (Novagen #69872) by PCR and NdeI-SalI digest of human GSK3α(1-97) from GST-3xVSVG-GSK3α(1-97) with the primers gcgccatATGGAGCAGAAACTCATCTCTGAAGAGGATCTGGAGCAGAAACTCATCTCTGAAGAGGATCTGGAGCAGAAACTCATCTCTGAAGAGGATCTGcctcccagtgcaggtgc (forward; capitals indicate 3xVSVG) and gcgcctcgagCCCTTGGAAGTATAGGTTCTCcagggtggtggaggggagc (reverse; capitals indicate a tobacco etch virus protease cleavage site). GST-3xHA-IκBα(1-62) was cloned into the BamHI and EcoRI sites of pGEX-4T-1 (3xHA) by PCR of human IκBα(1-62) from pINCY (Open Biosystems #IHS1380-97433845) with primers containing restriction sites for BamHI and EcoRI. 3xGluGlu-c-jun-His_6_ was cloned into the NcoI and SacI sites of pET-29b by PCR and NcoI-SacI digest of human c-jun from pBluescriptR (Open Biosystems #MHS6278-202808200) with the primers gcgcccATGGAGTATATGCCGATGGAGGAGTATATGCCGATGGAGGAGTATATGCCGATGGAGGGATCCGAATTCatgactgcaaagatg (forward; capitals indicate 3xGluGlu followed by BamHI and EcoRI restriction sites) and gcgcgagctcGAATTCGGATCCaaatgtttgcaactgctgcg (reverse; capitals indicate BamHI and EcoRI restriction sites added for future applications). The C-terminal hexahistidine tag was brought in frame by QuikChange II XL site-directed mutagenesis (Agilent #200521) using the following primers: catttggatccgaattcgcgagctccgtcgacaagc (forward) and gcttgtcgacggagctcgcgaattcggatccaaatg (reverse). GST-3xMyc-ERK2 was subcloned into pGEX-4T-1 (3xMyc) by digesting pGEX-4T-1 3xFLAG-ERK2 (Ref. ^95^) with BamHI and SalI to release rat ERK2 and ligating into similarly digested pGEX-4T-1 (3xMyc). GST-3xMyc-ERK2 was rendered catalytically inactive by a K52R mutation^96^ substituted through site-directed mutagenesis with primers gctcaaaaggactgattttcctgatagcaactcgaactttg (forward) and caaagttcgagttgctatcaggaaaatcagtccttttgagc (reverse) to yield GST-3xMyc-ERK2(K52R). GST-3xFLAG-HSP27 was cloned into the BamHI and EcoRI sites of pGEX-4T-1 (3xFLAG) by PCR of human HSP27 from pOTB7 (Open Biosystems #MHS6278-202829894) with primers containing restriction sites for BamHI and EcoRI. All plasmids will be available through Addgene.

### Bead conjugation

Five million MagPlex microspheres (Luminex) of a single bead region were added to a microcentrifuge tube (USA Scientific #1415-2500). Beads were pelleted at 8,000 rcf for 2 min and resuspended in 100 μL Milli-Q H2O. Beads were vortexed quickly and sonicated (Cole-Parmer #08849-00) for 20 sec to resuspend. Beads were pelleted again (8,000 rcf for 2 min) and resuspended by vortexing and sonication in 80 μL 100 mM monobasic sodium phosphate (pH 6.2). The bead surface was activated by adding 10 μL sulfo-NHS (50 mg/mL in H_2_O; Thermo #A39269) and 10 μL EDC (50 mg/mL in H_2_O; Thermo #77149). Beads were then gently mixed by vortexing and incubated at room temperature for 20 min with gentle vortex mixing every 10 min. Beads were pelleted (8,000 rcf for 2 min) and washed twice with 50 mM MES (pH 5.0). The activated beads were resuspended in 100 μL 50 mM MES (pH 5.0) and coupled with 10 μg carrier-free anti-epitope tag antibody (c-myc: Santa Cruz #sc-40; FLAG: Sigma #F1804; GluGlu: Biolegend #901801; HA: Roche #11867423001; VSVG: Sigma #SAB4200695) in a total volume of 500 μL 50 mM MES (pH 5.0). Beads were vortexed, wrapped in aluminum foil, and rocked at room temperature on a nutator for 2 hr. After conjugation, beads were pelleted and washed with 500 μL PBS + 0.1% BSA + 0.05% sodium azide, then twice washed with PBS + 0.02% Tween-20. The conjugated beads were resuspended in 250 μL PBS + 0.1% BSA + 0.05% sodium azide and counted on a Z1 Coulter counter (Beckman Coulter #6605698). Beads were stored wrapped in aluminum foil at 4°C in a dark box until use.

### Kinase assay

A mixture of antibodies for kinase immunoprecipitation (Table 1) was diluted in 50 μL 1% bovine serum albumin (BSA), 50 mM Tris-HCL (pH 7.5), 150 mM NaCl, 0.05% Triton X-100 (blocking buffer), added to each well of a Protein A/G coated strip plate (Pierce #15138), and rocked overnight at 4°C. Unbound antibody was aspirated and the plate washed three times with 200 μL blocking buffer. 75 μg of cellular extract in 100 μL kinase lysis buffer was added to the antibody coated wells and rocked at 4°C overnight. Lysate was aspirated and the plate washed twice with 200 μL 50 mM Tris (pH 7.5), 150 mM NaCl followed by an additional two washes with 30 mM Tris (pH 7.5), 22.5 mM MgCl_2_, 7.5 mM β-glycerophosphate, 1.5 mM EGTA, 300 μM sodium orthovanadate, 300 μM dithiothreitol (kinase wash buffer). The *in vitro* reactions were initiated with 60 μL of a three-part mixture containing 1) 20 μL kinase wash buffer; 2) 20 μL 30 mM Tris (pH 7.5), 45 mM MgCl_2_, 7.5 mM β-glycerophosphate, 1.5 mM EGTA, 300 μM sodium orthovanadate, 600 μM dithiothreitol, and 3 mM ATP added immediately before use; and 3) 20 μL substrates (Table 1) diluted in 1% BSA in Milli-Q H_2_O. Tubes for substrate dilution were pre-passivated for 30 min at room temperature in solution 1+3. The final *in vitro* kinase reactions were therefore 20 mM Tris (pH 7.5), 22.5 mM MgCl_2_, 5 mM β-glycerophosphate, 1 mM EGTA, 200 μM sodium orthovanadate, 300 μM dithiothreitol, 1 mM ATP, and 0.33% BSA. After initiation, the wells were sealed (Biorad #MSB1001) and the plate incubated for 1 hr at 37°C on a Jitterbug mixer (Boekel Scientific #130000) with mix setting set to 1. *In vitro* reactions were terminated with 60 μL ice-cold 20 mM EDTA. After mixing, samples were transferred to a storage plate (Abgene #AB-0859) and analyzed on a Luminex instrument or frozen at –80°C until detection.

For experiments involving IKK inhibition *in vitro*, SC-514 (10 μM; Sigma-Aldrich #SML0557) was added to the lysate before the overnight incubation on the immunoprecipitation plate. During the *in vitro* reaction, the concentration of ATP was reduced to 30 μM in solution 2, and 10 μM SC-514 was spiked into each well after the 60 μL three-part mixture was added.

### Luminex detection

Anti-epitope tag beads were mixed together at a concentration of 40,000 beads per tag/mL in 1% BSA, 20 mM Tris-HCl (pH 7.5), 137 mM NaCl (Luminex block buffer), and 50 μL of the bead mixture (~2000 beads per tag) was aliquoted to each well of a 96-well plate. All bead washes on the 96-well plate involved magnetizing on a magnetic plate (Millipore #40-285) for 2 min and forcefully inverting to remove buffer. The beads were washed twice with 20 mM Tris-HCl (pH 7.5), 137 mM NaCl, 0.05% Tween-20 (Luminex wash buffer). After removal of buffer, 20 μL of 20 mM Tris-HCl (pH 7.5), 1.24 M NaCl, 0.25% Tween-20 was added and then 30 μL of the kinase assay reaction to each well to enable epitope-tag immunoprecipitation to occur under high-salt conditions at a final concentration of ~0.5 M NaCl. The plate was placed on a plate shaker (IKA #0003208001), shaken at 1100 RPM for 30 sec, and incubated for 1 hr at 600 RPM. The plate was washed twice with 20 mM Tris-HCl (pH 7.5), 487 mM NaCl, 0.1% Tween-20 and then resuspended in 50 μL Luminex block buffer containing diluted anti-phosphosubstrate antibodies (Table 1). The plate was shaken at 1100 RPM for 30 sec and incubated for 2 hr at 600 RPM. After, the wells were washed twice with Luminex wash buffer, and the beads were resuspended in 50 μL Luminex block buffer containing a 1:10,000 dilution of biotinylated anti-rabbit antibody (Jackson Immunoresearch #111-066-144). The plate was shaken at 1100 RPM for 30 sec and incubated for 1 hr at 600 RPM. After, the plate was washed twice with Luminex wash buffer, and the beads were resuspended in 50 μL Luminex block buffer containing a 1:5,000 dilution of streptavidin-phycoerythrin (Prozyme #PJ31S). The plate was shaken at 1100 RPM for 30 sec and incubated for 30 min at 600 RPM. Finally, the plate was washed once with Luminex wash buffer, and beads were resuspended in 125 μL of Luminex block buffer for analysis. Plates were read on a MAGPIX (Millipore) and median fluorescence intensity for each bead region was recorded.

### Protein induction and purification

Plasmids were transformed into BL21-CodonPlus (DE3)-RIPL Competent Cells (Agilent #NC9122855), and single colonies were grown as overnight cultures in 5 mL terrific broth (TB) at 37°C with shaking at 250 rpm (New Brunswick Scientific #I26/I26R). The overnight culture was used to inoculate 250 mL TB in a beveled flask and continue growth at 37°C and 250 rpm. Upon reaching an optical density of 0.7–0.9, protein production was induced with 0.4 mM isopropyl β-D-1-thiogalactopyranoside (IPTG) (Sigma #I6758), and the culture was incubated at 37°C and 250 rpm for 4 hr. Cells were pelleted and mechanically lysed using an EmulsiFlex B15 (Avestin) at 60 psi as previously described^13^. Clarified extracts containing GST fusion proteins were passed through a 5 mL GSTrap column (GE Healthcare #17-5131-02) at 0.1 mL/min using an ÄKTAprime Plus chromatography system (GE Healthcare). The GSTrap column was washed with 20–30 mL of 25 mM sodium phosphate (pH 7.2), 150 mM NaCl. Bound proteins were cleaved off the column with thrombin diluted 1:250 in thrombin cleavage buffer (Novagen #69671-3). The thrombin solution was loaded manually into the column with a syringe, and the column was rocked overnight at 4°C. The column was eluted with 25 mM sodium phosphate (pH 7.2), 150 mM NaCl with an ÄKTAprime Plus operating at a flow rate of 0.1 mL/min. Fractions (3 mL) were collected and analyzed for protein content by SDS-PAGE and coomassie blue staining. Thrombin was removed from protein-containing fractions with 40 μL p-aminobenzamidine beads (Sigma #A7155) washed twice with 400 uL 137 mM NaCl, 2.7 KCl, 10 mM Na_2_HPO_4_, 1.8 mM KH_2_PO_4_. After rocking for 30 min at room temperature, the fractions were split into microcentrifuge tubes and beads were pelleted by centrifugation at 10,000 rcf (Beckman Coulter, Microfuge® 18). Supernatants were pooled, supplemented to 10% glycerol, and frozen at –80°C for storage.

Clarified extracts containing hexahistidine fusion proteins were passed through a 5 mL Ni-Sepharose (GE Healthcare #17-5268-01) packed column at 0.5 mL/min using an ÄKTAprime Plus chromatography system. The Ni-Sepharose column was washed with 20-30 mL of 25 mM sodium phosphate (pH 7.2), 250 mM NaCl, 20 mM imidazole. Protein was eluted by an elution buffer gradient of 25 mM sodium phosphate (pH 7.2), 250 mM NaCl, 20–500 mM imidazole and collected as 3 mL fractions from the ÄKTAprime Plus. Fractions were analyzed for protein content by SDS-PAGE and coomassie blue staining, supplemented to 10% glycerol, and frozen at –80°C for storage.

### Immunoblotting

500 μg of unstimulated HT29 lysate was immunoprecipitated onto Protein A/G coated wells (Pierce #15138) following the kinase assay protocol but with 0.5 μg of IKKα (Cell Signaling Technology #2682) or 0.5 μg of naïve rabbit IgG (Jackson Immunoresearch #011-000-002). Wells were washed twice with 200 μL 50 mM Tris (pH 7.5), 150 mM NaCl and twice with 200 μL 30 mM Tris (pH 7.5), 22.5 mM MgCl_2_, 7.5 mM β-glycerophosphate, 1.5 mM EGTA, 300 μM sodium orthovanadate, 300 μM dithiothreitol before eluting with 50 μL of 62.5 mM Tris-HCL (pH 6.8), 2% SDS, 10% glycerol, 0.01% bromophenol blue, 100 mM dithiothreitol and incubating at 100°C. Immunoblotting of immunoprecipitated proteins was performed as described^13^ with the following primary antibodies: IKKα (1:1000, Cell Signaling Technology #2682) and IKKβ (1:1000, Cell Signaling Technology #8943). Both species were detected with IrDye800-conjugated anti-rabbit secondary (Licor #926-32211).

## ACKNOWLEDGMENTS

We thank Karin Jensen for assistance with molecular cloning and early pilot experiments and Daniel Gioeli, Kristen Naegle, and the UVA Systems Biology Journal Club for reviewing the manuscript. This work was supported by the David & Lucile Packard Foundation #2009-34710 (K.A.J.), the National Institutes of Health #U01-CA215794 and #R21-AI105970 (K.A.J.), a Cardiovascular Research Training Grant Fellowship #T32-HL007284 (C.M.S.), a University of Virginia Vice-President of Research Fund for Excellence in Science and Technology award (K.A.J.) and 3 Cavaliers award (K.A.J.), a Translational Health Research Institute of Virginia seed collaboration research award (K.A.J.), and a UVA Cancer Center support grant #P30-CA044579.

## AUTHOR CONTRIBUTIONS

C.M.S. was responsible for conceptualization, investigation, formal analysis and drafting the manuscript. K.A.J. was responsible for conceptualization, formal analysis, reviewing and editing the manuscript, supervision, project administration, and funding acquisition.

---

**Supplemental Figure S1.**
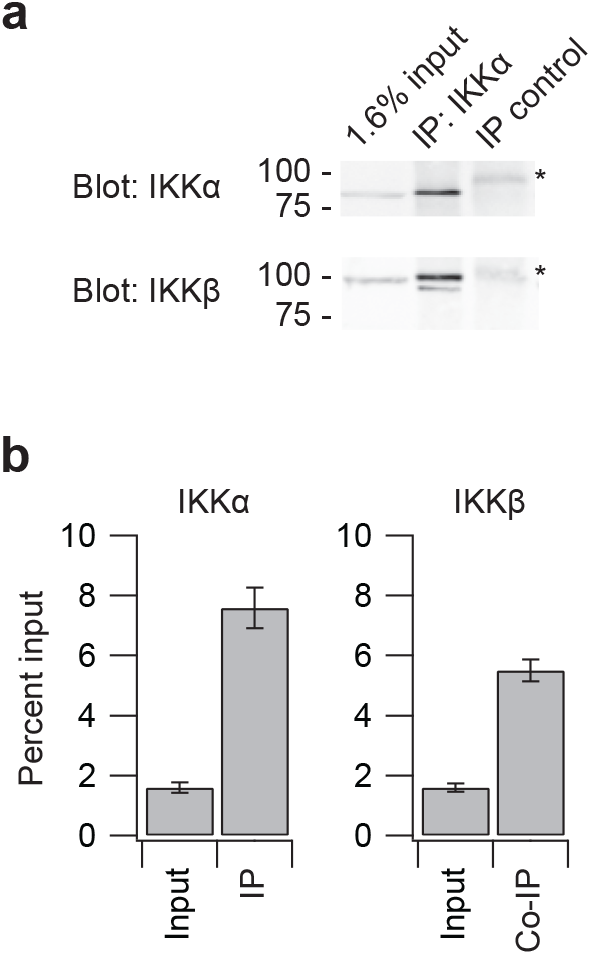
Co-immunoprecipitation of IKKβ in a microplate-based immunoprecipitation of IKKα. **(a)** Immunoblot of IKKα and IKKβ after anti-IKKα immunoprecipitation of 500 μg lysate on Protein A/G microplates. Immunoprecipitation (IP) control uses an equivalent amount of naïve rabbit IgG. Asterisk indicates a nonspecific band from incompletely reduced IgG heavy chain. **(b)** Quantification of IKKα and IKKβ in IKKα immunoprecipitates relative to input. Data are shown as the mean ± s.e.m. of *n* = 4 independent immunoprecipitations.

**Supplemental Figure S2.**
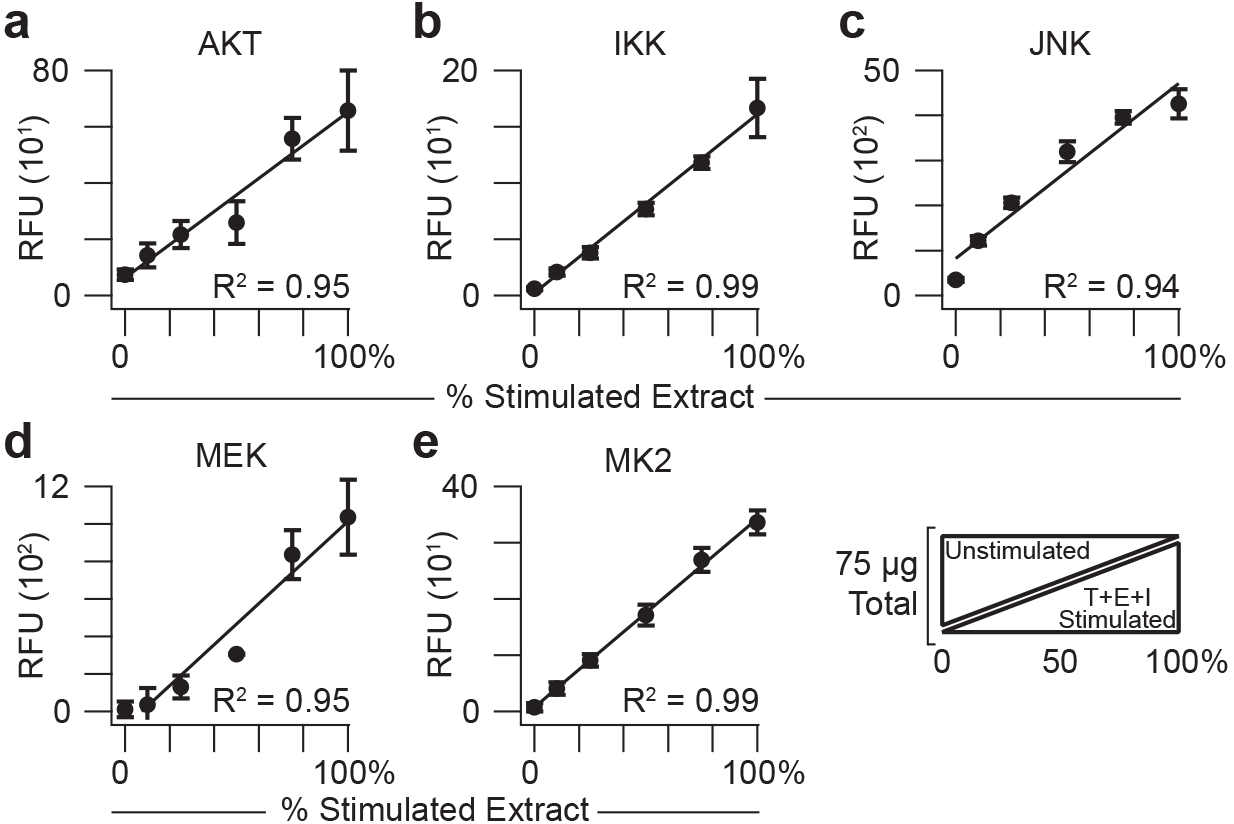
Response characteristics of the fiveplex kinase assay with titrated extracts from AC16-CAR cells. Relative *in vitro* phosphorylation of substrates for **(a)** Akt, **(b)** IKK, **(c)** JNK, **(d)** MEK, and **(e)** MK2 was measured together as a function of proportionately stimulated cell extract added. Data are shown as the mean relative fluorescence unit (RFU) ± s.d. of *n* = 4 assay replicates of 75 μg extract mixed in the indicated proportions of unstimulated AC16-CAR cells and cells treated with 20 ng/ml TNF, 100 ng/ml EGF, and 500 ng/ml insulin for 15 min (T+E+I stimulated).

**Supplemental Table S1.**
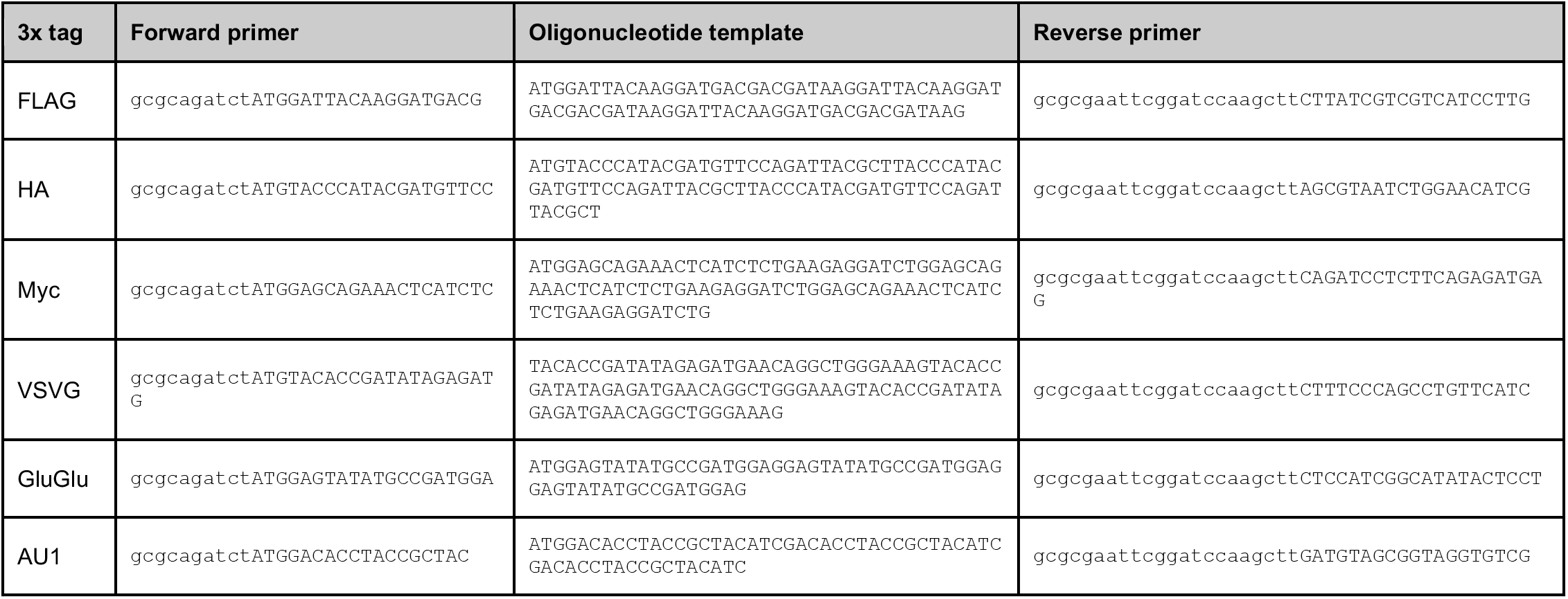
Oligonucleotides used for 3x epitope tag cloning. HindIII–BamHI–EcoRI restriction sites were added to the reverse primers for subsequent cloning of substrates.

**Supplementary Note.** Calculation of epitope-capture bead binding capacity at saturation.

The diameter of MagPlex microspheres is 5.6 μm, giving rise to 4π(5.6 μm/2)^2^ = 99 μm^2^ of surface area per bead. The average diameter of an IgG molecule is ~30 nm ^106^, whose maximum projected area on the bead surface is approximated as π(0.03 μm/2)^2^ = 0.0007 μm^2^. Taking the ratio of these two areas yields (99 μm^2^ per bead) / (0.0007 μm^2^ per IgG) = ~140,000 IgG per bead. Each anti-epitope IgG molecule is bivalent, and each substrate is trivalent because of its 3x epitope tag (see Methods). We assumed each substrate was avidly bound to the bead at two sites; thus, ~140,000 substrate molecules could be bound to each bead at saturation. The terminated kinase reaction is incubated with ~2000 beads per barcode, giving (~140,000 substrate molecules per bead) × (~2000 beads per reaction) / (6.02 × 10^23^ molecules per mole) = ~0.47 femtomole per reaction.

